# Recombulator-X: a fast and user-friendly tool for estimating X chromosome recombination rates in forensic genetics

**DOI:** 10.1101/2023.03.31.535050

**Authors:** Serena Aneli, Piero Fariselli, Elena Chierto, Carla Bini, Carlo Robino, Giovanni Birolo

## Abstract

**Background and Objective:** Genetic markers (especially short tandem repeats or STRs) located on the X chromosome are a valuable resource to solve complex kinship cases in forensic genetics in addition or alternatively to autosomal STRs. Groups of tightly linked markers are combined into haplotypes, thus increasing the discriminating power of tests. However, this approach requires precise knowledge of the recombination rates between adjacent markers.

Recombination rates vary across the human genome and cannot be automatically derived from linkage physical maps. The International Society of Forensic Genetics recommends that recombination rate estimation on the X chromosome is performed from pedigree genetic data while taking into account the confounding effect of mutations. However, the only existing implementations that satisfy these requirements have several drawbacks: they were never publicly released, they are very slow and/or need cluster-level hardware and strong computational expertise to use.

In order to address these key concerns, we developed Recombulator-X, a new open-source Python tool.

**Methods:** The most challenging issue, namely the running time, was addressed with dynamic programming techniques to greatly reduce the computational complexity of the algorithm, coupled with JIT compilation to further increase performance. We also extended the statistical framework from STR to any polymorphic marker.

**Results:** Compared to the previous methods, Recombulator-X reduces the estimation times from weeks or months to less than one hour for typical datasets. Moreover, the estimation process, including preprocessing, has been streamlined and packaged into a simple command-line tool that can be run on a normal PC.

Where previous approaches were limited to small panels of STR markers (up to 15), our tool can handle greater numbers (up to 100) of mixed STR and non-STR markers.

**Conclusions:** In the genetic forensic community, state-of-the-art estimation methods for X chromosome recombination rates have seen limited usage due to the technical hurdles posed by previous implementations. Recombulator-X makes the process much simpler, faster and accessible to researchers without a computational background, hopefully spurring increased adoption of best practices. Moreover, it extends the estimation framework to larger panels of genetic markers (not only STRs), allowing analyses of sequencing-based data.

## Introduction

The analyses of DNA profiles for personal identification and kinship in forensic casework strongly rely on the biostatistical evaluation of the evidential weight under alternative mutually exclusive scenarios, whose specific probabilities are then combined into likelihood ratios (LRs). Short tandem repeats (STR) have been the markers of choice for such analyses due to their high discriminating capacity and genotyping ease using standard capillary electrophoresis typing techniques [1]. While autosomal DNA polymorphisms are most widely used in forensic practice thanks to their higher informativeness, particular caseworks require complementary information from other genomic regions including haploid markers. For instance, mitochondrial DNA is crucial when ancient or degraded genetic material is involved, while Y-chromosomal STRs are fundamental for the interpretation of mixtures involving a high ratio of female:male contribution [2]. Thanks to its unique features, halfway between autosomes and uniparental markers, the STR markers on the X chromosome (X-STRs) play a relevant role in challenging kinship testing, such as when the DNA from one of the parents is unavailable (kinship deficiency cases), half-sister or incest cases (Supplementary Figure 1, for details, see Supplementary Materials). Moreover, in paternity analyses with inconclusive or statistically weak results, for instance in case of genetic inconsistencies or poor amplification from exhumed remains, adding X-STR markers can help in reaching an informative solution [3–7]. Over the last decade, use of sequencing-based techniques has become increasingly widespread in forensic genetics [8–11]. This brought back interest in other types of markers, especially SNPs, which can be analysed either in combination with STRs or alone [12–17]. Whilst more SNPs are necessary to reach the discrimination capacity of STRs, the possibility of genotyping large number of markers simultaneously from low quantities of input DNA or from degraded material has opened the floodgates to new forensic applications also based on SNPs, such as ancestry inference, DNA phenotyping and investigative genetic genealogy, the latter being specifically designed on dense SNP data [18–24]. Nevertheless, due to the almost exclusive attention traditionally given to STRs in forensics, most available tools do not support non-STR markers.

Forensic markers are located in the non-pseudoautosomal region of the X chromosome. This means that while females have two haplotypes, one inherited from the mother (the maternal haplotype) and one from the father (the paternal haplotype), males have just a single haplotype inherited from the mother. This haplotype is a mixture of the mother’s two haplotypes as a result of the recombination process. Since the genetic size of the X chromosome is about 155Mb [25] and assuming a 50Mb physical distance between markers to ensure independence, a maximum of 3-4 markers can be simultaneously analysed as independently segregating. For this reason, traditional analyses of highly polymorphic haplotypes, consisting of X-STR markers organised into “linkage groups” or “clusters”, were devised in order to increase the evidential weight, which would be otherwise statistically inconclusive [6]. Nowadays four different X-STR linkage groups are routinely used for forensic applications [3,26–44]. However, it has been shown that, while well spaced along the X chromosome, some of these linkage groups cannot be considered truly independent from each other [45–47]. The consequent violation of the independence assumption requires proper considerations in the biostatistical evaluation of kinship. Moreover, although it was originally assumed that recombination did not happen within linkage groups, later studies have demonstrated that, albeit rare, recombination may occur, thus motivating the evaluation of recombination rates for markers both between and within the same cluster [6,48,49]. Indeed, the latest recommendations of the International Society for Forensic Genetics (ISFG) about the use of X-STRs in kinship analyses clearly indicate the precise knowledge of recombination rates between markers included in in-house and commercial X-chromosomal multiplex PCR assays as a prerequisite to unbiased estimates of kinship likelihood ratios (LRs) [6]. The available software for kinship LR calculations, with FamLinkX being the most widely used, infers neither recombination nor mutation rates, which are instead expected to be known a priori [45,46]. However, the evaluation of such measures is not straightforward and may appear computationally intensive. As also highlighted by recent works on the use of the X chromosome in forensics [3,50], the analytical and statistical issues deriving from genetic linkage and the lack of software addressing such issues are actually hindering the proper applications by leading to significant biases in the quantification of the genetic evidence. For this reason, technical advancements in this field are highly encouraged [3,6,50].

Recombination rates are known to vary across the human genome and cannot be automatically derived from combined linkage physical maps [51]. In the case of forensic X-STRs, recombination rates have been either inferred from population samples through high-density multi-point single nucleotide polymorphism (SNP) data [52] or directly estimated in large pedigree-based studies [44,48,49,53]. However, while population-based approaches may suffer from long-term population size changes and selection effects, pedigree studies infer recombination across a few generations by directly observing the inheritance of alleles from parents to offspring [54]. Indeed, the ISFG’s guidelines recommend that recombination rates should be primarily estimated from family-based studies [6]. For these reasons, further pedigree-based studies are expected in the future to comply with the steady increase in the number of X-chromosomal markers described for forensic applications [3] and to investigate possible population-specific variability in recombination rates [55].

Recombination between X-chromosomal markers only happens in female meiosis. This entails that only females can provide information on recombination events, while haploid males can be used to phase their mother/offspring. Such events are more easily observed between mother and sons, since genotyping sons immediately yields the recombined maternal haplotypes. Ideal linkage-informative families in pedigree studies are therefore three-generation families, including maternal grandfather, mother and one or more sons. In such families, labelled as *type I*, the mother can be phased using the grandfather and thus recombination events between the maternal haplotypes can be directly observed in the sons. Also informative are two-generation families consisting of one mother and two or more sons, labelled as *type II* (Figure 1A). Here the maternal haplotypes cannot be determined given the lack of the grandfather. Hence the need for multiple sons to be evaluated together in order to discern, among all possible maternal phasings, those that can better explain multiple recombination events and thus also give information about recombination rates. Moreover, not only sons can be used: when the father genotype/haplotype is available, the maternal haplotype can also be retrieved from a daughter after phasing (Figure 1B).

**Figure 1.**
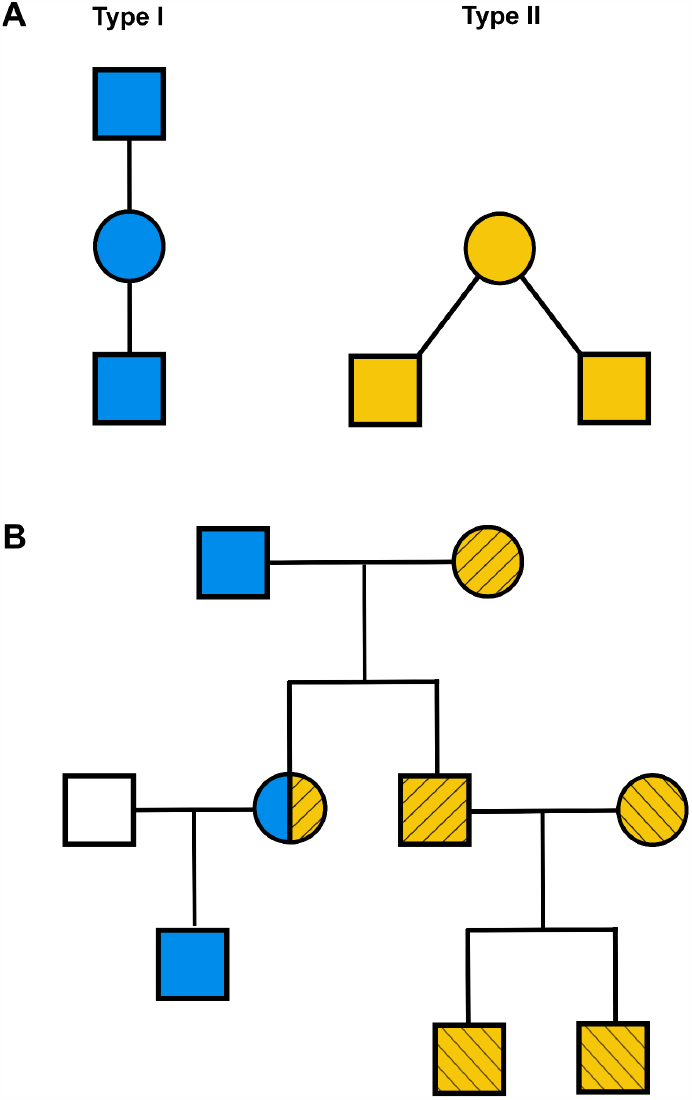
A) Three and two-generation family configurations useful for inferring recombination events (type I in blue on the left and type II in yellow on the right). B) An example pedigree in which one type I (in blue) and two type II families (in yellow with stripe patterns) can be extracted.

The standard statistical approach for the estimation of recombination rates from pedigrees computes the likelihood of kinship by taking into account all possible recombinations within the maternal haplotype, resulting in the exponential complexity of the original algorithm [48]. Despite a new implementation in C++ that allows multi-core parallelization, this approach remains too slow to handle panels of more than 15 X-STRs [49]. Such limitation clashes with the increasing capability of forensic laboratories to simultaneously investigate larger panels of DNA markers favoured by advances in standard capillary electrophoresis typing techniques and the growing use of massively parallel sequencing (MPS) technology [8–17].

Moreover, the implementations of the estimation algorithm were never released to the public, even though they are available upon request from the authors (who kindly provided us with the original R script). They also do not include necessary steps such as data parsing and preprocessing, requiring some R programming knowledge from the user. All of these issues make performing the estimation on new datasets quite onerous, to the point that some recent studies resorted to less accurate but simpler approaches that can be solved manually [56–59].

In order to make the estimation of recombination rates between X chromosomal markers faster and more accessible, we developed the first open-source, batteries-included software with optimised algorithms that allows the user to perform the estimation from a standard pedigree file in just one command. The new algorithms implement the same statistical framework of the previous work [48], without approximations or limiting assumptions, but extending its applicability also to other types of polymorphisms (e.g., SNPs and INDELs). Taking advantage of dynamic programming and other optimization techniques, we were able to drastically reduce computational time, also allowing us to handle an increased number of markers than previously possible. We released this work as a Python module named “Recombulator-X”, which is the first open-source software for the estimation of the recombination and mutation rates for all types of genetic markers (Figure 2). Beyond the optimised implementations of the estimation method, it includes a command-line tool (requiring no programming knowledge), extensive documentation and usage examples, all available in a GitHub repository (https://github.com/serena-aneli/recombulator-x) and a dedicated website (https://serena-aneli.github.io/recombulator-x/).

**Figure 2.**
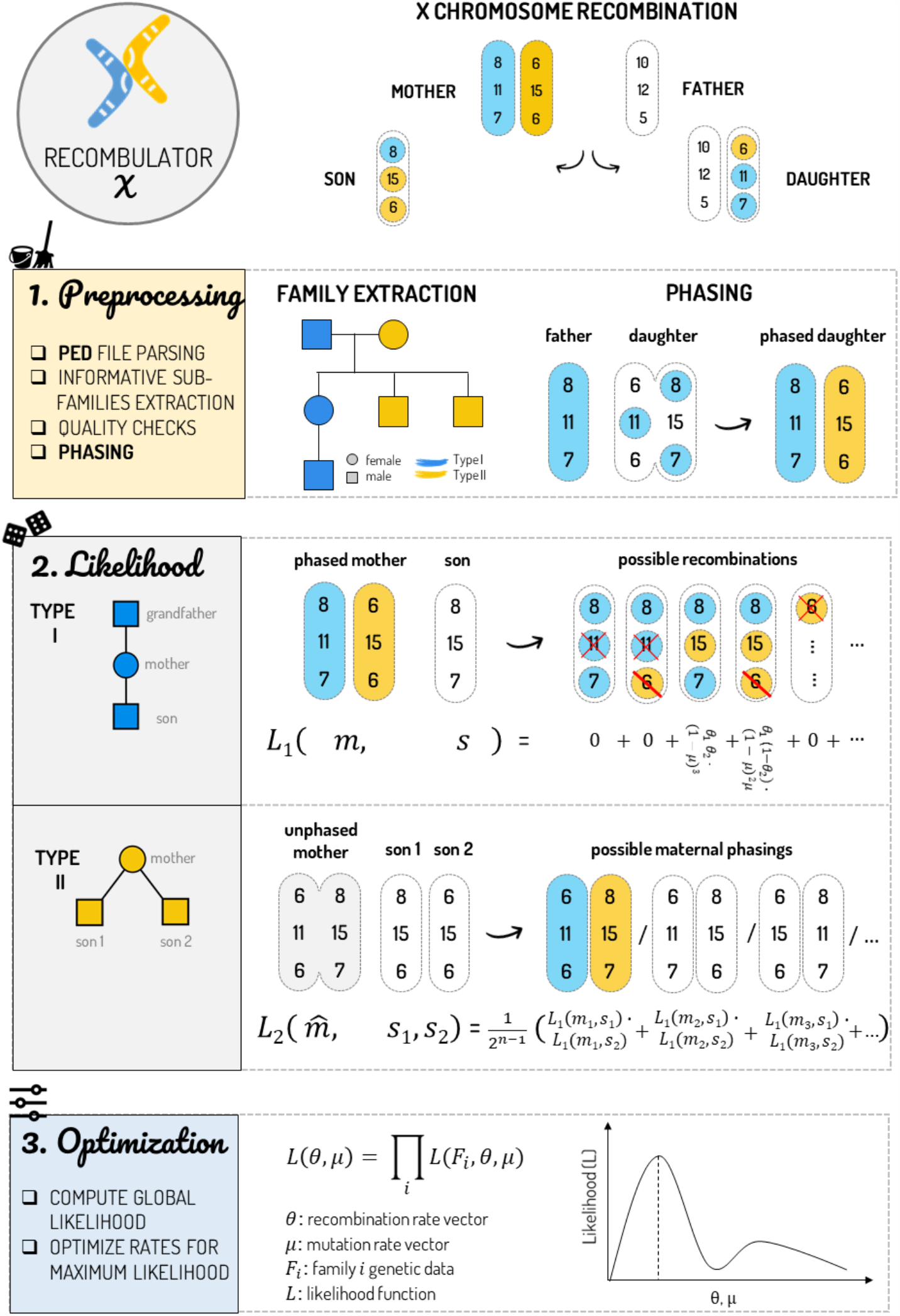
The steps of Recombulator-X are represented on the left column of the figure, while we reported a simplified example on the right using just three X-STRs. 1) Preprocessing: Recombulator-X reads the PED file, performs preliminary quality checks, extracts the informative type I and type II families and phases all the females, whenever their father is available. 2) Likelihood computation, depending on the family type: in the case of a phased mother (type I family), the likelihood (L_1_) of each possible recombination is computed and summed up. Here, the red crosses indicate genetic incompatibilities (mutations greater than one repeat), while the single red lines correspond to compatible single-step mutations. When the grandfather is not available (and thus the mother cannot be phased, type II family), this process is repeated for each possible maternal phase (L_2_). 3) In the last step - optimization - the likelihood of the entire dataset is computed by multiplying together the likelihood of each family and Recombulator-X searches the parameters (recombination and mutation rates) that maximize the global likelihood.

## Methods

Recombulator-X follows the general statistical framework and estimation strategy introduced in [48]. There, the authors define a likelihood function that computes the exact probability of observing a pedigree, given recombination and mutation rates as parameters. Then, they use standard optimisation techniques (the L-BFGS-B method implemented in the *optim* function in R) to find those rates that maximise the likelihood of the dataset. Our main contribution is introducing a much faster implementation of the likelihood function (based on an optimised algorithm) which yields the exact same probability as the original, that is, without resorting to approximation.

### Statistical framework and algorithmic optimization

We present here the statistical framework first introduced in [48] and show the parts that resulted amenable to optimization.

A haplotype of STR markers can be described as a vector *x* = (*x*_1_, …, *x*_*n*_) of positive rational numbers (repeats can be fractionary). Let *m* = (*m*^1^, *m*^2^) be the mother’s haplotypes and be the child’s maternal haplotype (the one inherited from the mother through recombination of her haplotypes). Possible recombinations are represented by the *inheritance vector υ =* (*υ*_1_, …, *υ*_*n*−1_) where *υ*_*i*_ ∈ {1, 2} for all *i* (Supplementary Figure 2).

Other parameters are the recombination and mutation rate vectors *θ* and *μ* of length *n* − 1 and *n*, respectively. Rates are all in the 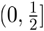 intervals.

Then we can define the likelihood of observing a child from a mother by a specific inheritance vector as follows:

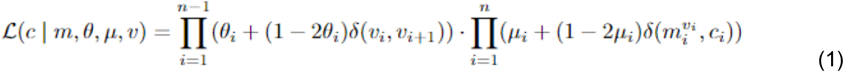

where *δ*(*a, b*) is 1 when *a* = *b* and 0 otherwise.

Then, by summing over all possible 2^*n*^ inheritance vectors in *V* = {1, 2}^*n*^, we obtain the likelihood of a child’s haplotype: *V* = {1, 2}^*n*^, we obtain the likelihood of a child’s haplotype:

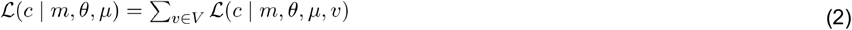

A son’s maternal haplotype can be observed directly from his genotype. For daughters, it can be inferred by subtraction of the father haplotype, when it is available.

This definition of likelihood is enough for type I families (Figure 1A), where the mother’s haplotypes can be determined using the grandfather’s haplotype and multiple children are handled as independent recombination events, thus multiplying their likelihood together:

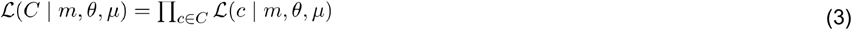

where *C* is the set of the children’s haplotypes.

For type II families, where only the mother’s genotype is known but not her haplotypes, equation 3 must be extended by further conditioning on the set *M* of all possible mother’s haplotypes given her known genotype as follows:

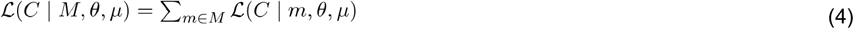

The original exponential-time algorithm, which we will call the *direct* algorithm, iterates over all possible *2*^*n*^ recombinations of *n* markers, following directly equation (2). However, for any two inheritance vectors that are equal up to marker *i*, it can be seen that, in equation (1), all products up to maker *i-1* are the same. These repeated sub-computations can be avoided with dynamic programming techniques and indeed by factoring them out we obtained an optimised linear time-complexity algorithm for computing (2) and (3) with no loss of precision, which we will call the *dynamic* algorithm. This is the case for type I families, where the mother’s haplotypes are known. For type II families, where the grandfather is unavailable and only the mother’s genotype is known, an additional iteration on all the possible *2*^*(n-1)*^ maternal haplotypes is still needed as in equation (4). Thus, type II families require exponential time also with the dynamic algorithm, albeit going from *O(2*^*(2n-1)*^) of the original direct algorithm to *O(n*2*^*(n-1)*^). Our solution still provides an exponential speed-up with respect to the standard implementation (speed up is equal to *O*(2^*n*^/*n*)).

The new likelihood function is then used to estimate the recombination and mutation rates in the same way as in the original paper, by finding a minimum of the negative log-likelihood of the dataset with the rates as parameters, employing the L-BFGS-B method for bound constraints (the rates must be positive and smaller than 0.5) as implemented in the *scipy*.*optimize*.*minimize* function in the SciPy Python package.

### Likelihood implementations

Beyond the algorithmic improvements, other optimization techniques were explored to further reduce computation times. As a result, Recombulator-X includes multiple implementations of the likelihood computation, both of the original direct algorithm and the improved dynamic one: the *direct-loop, direct-numpy, dynamic* and *dynamic-numba* implementations.

The *direct-loop* is a straightforward implementation of the direct algorithm using loops, similar to the original R implementation. This version of the likelihood is arguably the simplest to understand and was thus used as a reference for testing the correctness of the more complex optimised versions. The same computation was also implemented using the fast vectorized operations offered by the NumPy package, with the label *direct-numpy*. However, this implementation is still exponential in time and also in space, since it requires intermediate results to be stored as large multidimensional arrays.

The dynamic programming algorithm is much faster with its linear complexity (for type I families), even when written using Python loops as in the *dynamic* implementation. However, it still benefits from being JIT compiled with the Numba package as the *dynamic-numba* implementation [60]. For type II families, to ameliorate the still exponential complexity, we introduced a further optimization, branching through the possible maternal phasings and computing partial likelihoods up to a certain marker, sharing part of the computations and discarding the branches with zero likelihood early.

Testing and benchmarking were performed by simulating random pedigrees from given recombination and mutation rates using generative functions (included in Recombulator-X). All benchmarks were averaged across ten runs on a workstation with an i9-12900F processor.

### Extension to non-STR markers

We extend the statistical framework from [48] to handle panels of arbitrarily mixed STR and non-STR polymorphisms. Single base substitutions are expected to be represented as single-letter codes, but generic strings are accepted to accommodate for more complex non-STR polymorphisms like INDELs. Internally we extend the numeric representation of alleles by encoding unique non-STR alleles with decreasing negative integers. In the likelihood definition, we keep the recombination part unchanged since it is not affected by the type of marker, but we need to extend the mutation part. So we replace equation (1) with the following:

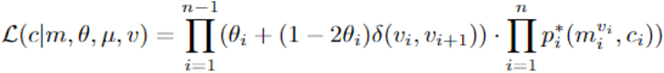

where 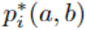 is the probability of mutation from an allele *a* to an allele *b*, which is defined differently depending on the type of marker. For STR markers, we define:

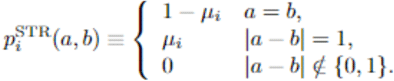

so that non-unit mutations (insertion or deletions of multiple or partial repeats) have zero probability since they are much less frequent than unit mutations. Instead for non-STR polymorphic markers, we define:

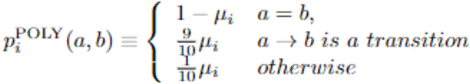

where transitions are single base mutations between purines (A and G) or pyrimidines (C and T), which are much more frequent than other single base substitutions or more complex polymorphisms [61].

### Additional features

While the dynamic programming algorithm for the likelihood function is arguably the core of the package, Recombulator-X original contributions also include non-trivial dataset parsing and preprocessing functions. Thanks to those, datasets can be read from the standard PED format as used by the PLINK software [62], a simple textual format that encodes both arbitrary relatedness and genetic information. Preprocessing functions then 1) build an arbitrarily complex graph for each group of related individuals (Supplementary Figure 3); 2) extract all subgraphs that can be used as type I or II families; 3) phase mothers and daughters whenever possible and finally yield the processed dataset ready for the estimation. During this preprocessing some consistency checks take place, alerting the user of eventual problems with the data. All these steps are wrapped into a single command line tool, that takes a pedigree file as input and outputs the recombination and optionally the mutation rates. This tool allows the user to run the entire estimation without programming knowledge.

## Results

In order to assess the reduction in the likelihood computation time and how it improves the whole recombination and mutation rate estimation process, we performed a series of benchmarks using simulated datasets with incremental number of markers.

The first benchmark compares the different algorithms and implementations of the likelihood function that are included in Recombulator-X. To see how the computation time is affected by the number of markers, we measured the average time to compute the likelihood of type I or type II simulated families in Figure 3. The exponential complexity of the direct algorithm is clearly visible, both for type I and type II families. The dynamic programming algorithm shows instead its linear complexity for type I families, allowing a virtually unlimited number of markers. Unfortunately, type II families remain problematic, even though the dynamic programming numba-optimised version is able to handle ten more markers than the numpy-vectorized direct implementation in the same time (from 11 to 21 markers).

**Figure 3.**
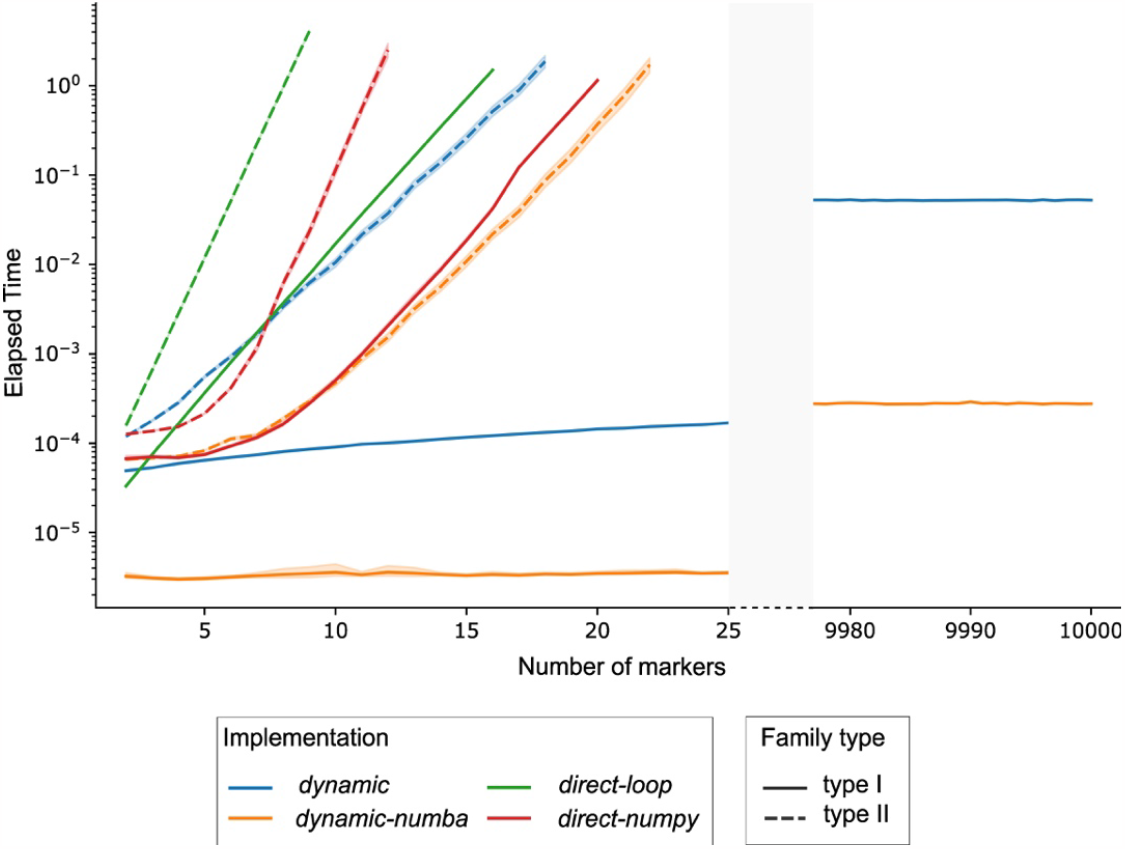
Mean time needed to compute the likelihood for one family typed over up to 10,000 markers. Each implementation is represented with a different colour, while the linestyle refers to family types. The y axis is in log scale. For each implementation, the number of markers was progressively increased until the computation time went above one second per family.

After testing the likelihood function implementations, we benchmarked the entire optimization procedure using the fastest implementation, dynamic-numba, in order to see how the improvements to the likelihood computations impact the whole process of recombination and mutation rate estimation. The results are reported in Figure 4.

**Figure 4:**
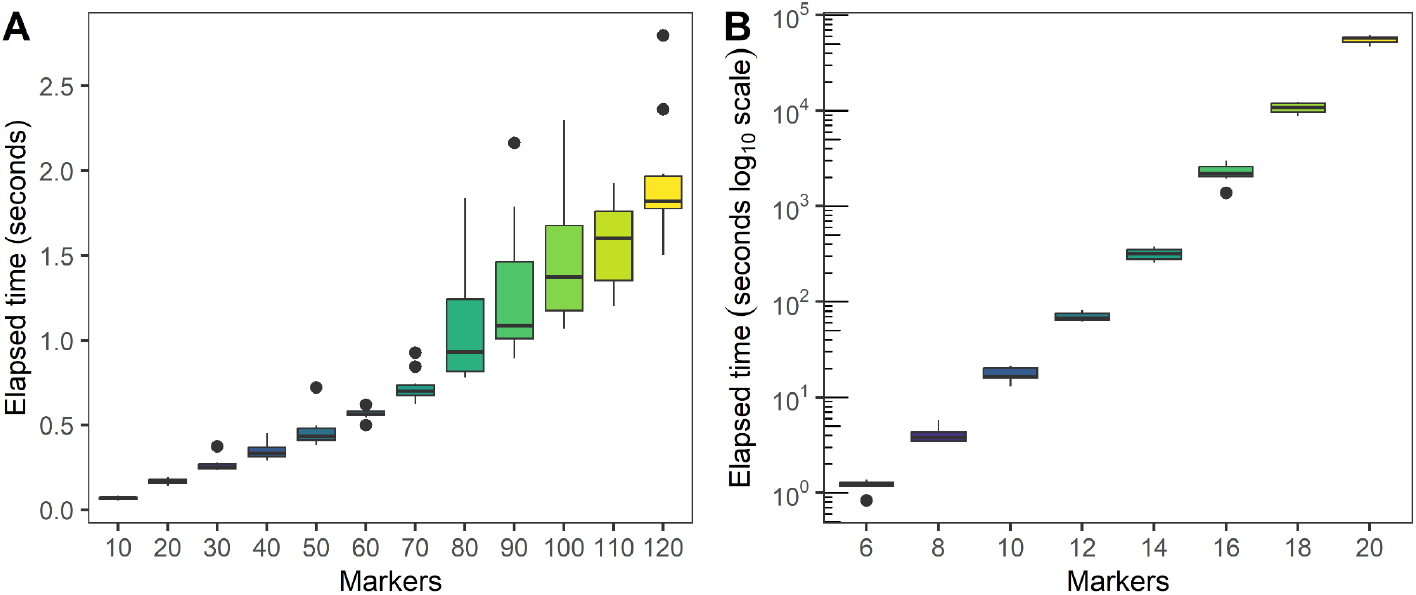
Recombination and mutation estimation times using the fastest (dynamic-numba) likelihood implementation depending on the number of markers. A simulated dataset of 100 type I families (A) and one of 100 type I and 100 type II families (B) were tested. Times are in seconds, with a logarithmic axis for the right panel.

Unsurprisingly, when considering only type I families, the estimation is very fast. For 100 families, the estimation of up to 100 markers is almost instantaneous, taking less than two seconds. However, at around 130 markers, we start having issues where the optimization fails to converge. Even raising the iteration limit for convergence and trying the other optimization methods available in SciPy did not allow the estimation to converge.

When also type II families are involved, the exponential complexity of the likelihood function poses a hard limit to the number of markers. For a dataset of 100 type I and 100 type II families, the computation times are much longer, roughly doubling for each additional marker (as expected). We stopped testing at 20 markers, where the average computational time was 16 hours.

While a direct comparison of Recombulator-X with previous software is difficult (the genetic data from the two previous studies are not publicly available), we also simulated datasets with the same size as in previous studies to have a comparison in more realistic scenarios. Nothnagel and colleagues analysed 216 type I and 185 type II families genotyped with a panel of 12 markers [48]. While they did not report the time required, they felt the need to say that a faster implementation was needed. Our method only takes 3.5 minutes on a simulated dataset with the same number of families. In a later analysis, involving 54 type I and 104 type II families with a panel of 15 markers, the authors developed a much faster parallel version in C++. However, due to the increased number of markers and the exponential complexity, the estimation process took a few months on a highly parallel computing system [49]. Our method takes 20 minutes to carry on the same task. While these times are not completely comparable given that they ran on different datasets, on different hardware and in different languages, it is clear that such a decrease in time cannot be attributed to those factors alone and that Recombulator-X, even with its current limitations, brings a substantial improvement over previous methods.

## Discussion

The proper biostatistical evaluation of the evidential weight in personal identification and kinship tests when dealing with X chromosome markers is a nagging problem in forensics, due to physical linkage [3,6,46,50,63]. Despite being crucial for unbiased formulations of the evidential weight, as also highlighted by the International Society of Forensic Genetics [6], few biostatistical tools for the evaluation of recombination rates between adjacent forensic markers along the X chromosome are available today.

Routine kinship analyses rely almost exclusively on commercial kits, such as the commonly used Argus X-12 QS which consists of 12 X-STRs [3,26–40,42–44]. However, current implementations of state-of-the-art statistical framework for estimation from pedigrees, besides being quite onerous to use, are already very slow for 12 markers and so unsuitable for larger panels without the availability of large computational resources [48,49]. Consequently, many recent works have been limited to a “manual” evaluation of the recombination rate which does not consider the mutation probability [56–59].

The growing use of next-generation sequencing technologies in the forensic fields, with the possibility of combining thousands of markers together, requires the development of new biostatistical frameworks scalable to a higher number of genetic markers [9]. Moreover, many commercially available NGS-based kits allow to combine STRs and other non-traditional markers, such as SNPs or INDELs [12–17]. In particular, SNPs have been increasingly appealing thanks to their technical features and informational power: their smaller amplicon size is crucial with samples of low quantity and poor quality (this is relevant since the majority of forensic analyses involves degraded DNA) [64] and they provide insight for predicting human appearance and the biogeographical origin of unknown sample donors or deceased/missing persons [65,66], thus ultimately resulting in new investigative leads. Additionally, given their lower mutation rate when compared to STRs, they were shown to be helpful in solving kinship cases [67,68]. Notably, the latest application of SNPs is investigative genetic genealogy where dense SNP data are jointly analysed to infer distant relationships (which in forensics indicate relatedness exceeding that of first cousins) [69]. For these reasons, an increasing number of commercial NGS-based kits have included X chromosomal SNPs and/or STRs to address complex kinship scenarios [14,18,70–76]. Nevertheless, complex kinship cases relying on many and mixed types of X chromosomal genetic markers cannot be addressed using the previous implementations for the inference of recombination rates, which are used, albeit with limitations, for STR markers.

In order to overcome these issues, we developed Recombulator-X, the first open-source tool for rapidly inferring X chromosome recombination rates. Our optimised algorithm is substantially faster than existing gold-standard methods, with no loss of accuracy since it is based on the same statistical framework. Performing the estimation on standard panels of 12 markers on a new dataset can now be done in minutes instead of days or weeks on a single PC. This will also enable new studies to experiment with larger panels than previously possible, going from a practical limit of around 15 markers to more than 25 for general datasets and one hundred when considering only type I families. Moreover, the extension to mixed STR and non-STR markers is especially relevant to enable sequencing-based panels.

No less important from a practical point of view is that the full implementation and source code (including dataset parsing and preprocessing) are available as a Python package. The repository also includes documentation, usage examples and a command line tool, greatly simplifying the estimation process for a non-technical user.

For all these reasons, we hope that Recombulator-X might transform the estimation of recombination rates from an arduous process requiring specialised expertise and hardware to a routine computational analysis that anyone can perform.

## Acknowledgements

We would like to thank Prof. Michael Nothnagel for kindly providing us with his original code and answering our questions. We also thank Programma Nazionale della Ricerca PNR 2021-2027 e PON “Ricerca e Innovazione” 2014-2020 – progetti di ricerca su tematiche “Innovazione” e “Green” (S.A.) and PNRR M4C2 HPC – 1.4 “CENTRI NAZIONALI”-Spoke 8 for fellowship support.” (P.F).

## Author Contributions

CR and CB introduced the problem to SA and GB and prompted the initial development. PF suggested the dynamic programming approach and provided much needed advice. SA and GB implemented the software, wrote its documentation and drafted the manuscript with input from all authors. EC, CB and CR provided critical feedback about the practical needs and issues in the forensics community. All authors revised and approved the final version of the manuscript.

## Data availability

Recombulator-X is freely available at https://github.com/serena-aneli/recombulator-x. Full documentation can be found on a dedicated website at https://serena-aneli.github.io/recombulator-x/.

