## Supplementary Materials for "Recombulator-X: a fast and user-friendly tool for estimating X chromosome recombination rates in forensic genetics"

**1 X chromosomal markers in forensics - an overview 2**

### 1 X chromosomal markers in forensics - an overview

The X chromosome is unique in the human genome. Indeed, the peculiar structure, as well as its mode of inheritance and functions, which are not shared by the Y chromosome and neither are by any of the autosomes, make it a fundamental tool for understanding gender-specific genetic differences, delving into the molecular basis of most Mendelian diseases, addressing the history of the human population, as well as solving complex kinship cases.

As highlighted by Balaton and colleagues in their paper "The eXceptional nature of the X chromosome", many characteristics differentiate the X chromosome from the rest of the genome (Balaton *et al.,* 2018). To begin with, the X chromosome is present in a single copy in males, while females have two copies, thus maintaining the original autosomal configuration. However, the exclusive inheritance mode, which depends on gender, is the most peculiar feature of the X chromosome. In males, the single copy of the X chromosome is transmitted as a single unbroken DNA chunk to the females of the next generation, while in females the two chromosomal copies recombine, in the same manner as the autosomes, thus increasing genetic variation across generations. One of the two reshuffled chromosomes is then passed to both male and female descendants. This mechanism makes the male copy of the X chromosome easier to study than the autosomes, because, in the same manner as the mitochondrial DNA (mtDNA) and the Y chromosome, the male copy can be considered an unbroken haplotype. Indeed, both the male X chromosome and the uniparental markers have been extensively used to reconstruct the past demographic events experienced by human populations. Actually, a certain degree of recombination is allowed even in males in order to enable proper chromosomal segregation during meiosis. The shuffling occurs exclusively at the chromosomal tips, the two homologous sub-telomeric regions known as pseudo-autosomal regions (PAR).

Although different types of genetic markers have been extensively used for personal identification and kinship analyses, such as Single Nucleotide Polymorphisms (SNPs) and Insertions/Deletions (INDELs), Short Tandem Repeats (STRs) are the preferred ones in forensic applications. STRs are genetic markers composed of 2 to 7 base pair long repeat units. The three characteristics that make the STRs widely used in forensics are: i) since they are highly polymorphic, their discriminating capacity between individuals is higher than both SNPs and INDELs, meaning that fewer markers are needed for identification purposes; ii) PCR-based technologies and automated capillary electrophoresis can be coupled in a straightforward analytical workflow to detect them; iii) a generally short amplicon length makes them useful when dealing with degraded DNA (Gomes *et al.,* 2020; Garcia *et al.,* 2022).

Specific combinations of X-STRs are included in commercial kits available on the market. The first one, composed of four X-STRs (DXS8378, DXS7132, HPRTB, and DXS7423), was the Argus X-UL from Biotype (Dresden, Germany), which was then updated with four additional markers (Argus X-8). The most recent and widely used kit is the Argus X-12 (Qiagen, Hilden, Germany; the Argus X-12 QS is its optimized newest version, which contains the same markers). Argus X-12 contains 12 X-STRs distributed along the X chromosome into four different linkage groups (see later on): LG1, DXS10148/DXS10135/DXS8378; LG2, DXS7132/ DXS10079/ DXS10074; LG3, DXS10103/ HPRTB/ DXS10101; and LG4, DXS10146/ DXS10134/ DXS7423.

While routinary casework is normally addressed with STRs, thanks to the technical advantages of NGS-based approaches which allow the co-analysis of STRs/SNPs panels while minimizing waste of sample, there has been a strong revival of interest in SNPs. Indeed, while more SNPs are necessary to reach the same informativeness as STRs, some peculiar features make them an interesting asset for particular forensic applications, such as DNA mixture interpretation, DNA phenotyping and personal identification/ kinship analyses with degraded DNA.

#### 1.1 Forensic applications

While standard identification and kinship cases (duos and trios) can usually be addressed by autosomal markers, the peculiar inheritance mode of X chromosomal markers makes them more informative than autosomal, Y-chromosomal and mitochondrial DNA markers, depending on the examined genetic relationship. Indeed, specific casework may greatly benefit from the inclusion or even the sole use of X chromosomal markers (Figure S1). A thoughtful general framework for identifying situations where the X chromosomal markers are useful in kinship analyses is reported in Pinto *et al.,* 2011 and Pinto *et al.,* 2012. Obviously, we may certainly exclude those situations involving a link "father-son" in both the main and the alternative hypotheses. For instance, a "paternal grandfather - granddaughter" vs "unrelated" case can not be solved by X chromosomal markers, given that the putative grandfather did not transmit his X chromosome to his son, whose sole X chromosome derives from his mother. Conversely, in paternity analyses with inconclusive or statistically weak results, such as when few genetic inconsistencies arise or in case of poor amplification results from exhumed remains, adding X-STR markers may help in reaching an informative solution. For instance, the finding of a few inconsistencies between the alleged father and the daughter may be explained by the occurrence of mutations or by a close relationship between the alleged and the real father. However, this relationship can be solved with X chromosomal markers: indeed, if the biological father is her grandfather or her half-brother, they will not share X chromosomal markers identical by descent (Tillmar *et al.,* 2017).

Moreover, in particular kinship cases, the inclusion/exclusion power inferable from X chromosomal markers is higher than with autosomes. For example, in half-sisters or deficiency paternity cases, where a father-daughter hypothesis is considered, the usage of X-STRs may be helpful given that there is a 100% transmission of the father's X chromosome to his daughter. Another example regards, as above-referred, incest: when the investigation concerns whether the victim's father or the victim's brother has fathered the victim's daughter, autosomal markers, given the genetic relatedness between the individuals involved in testing, are generally not so helpful.

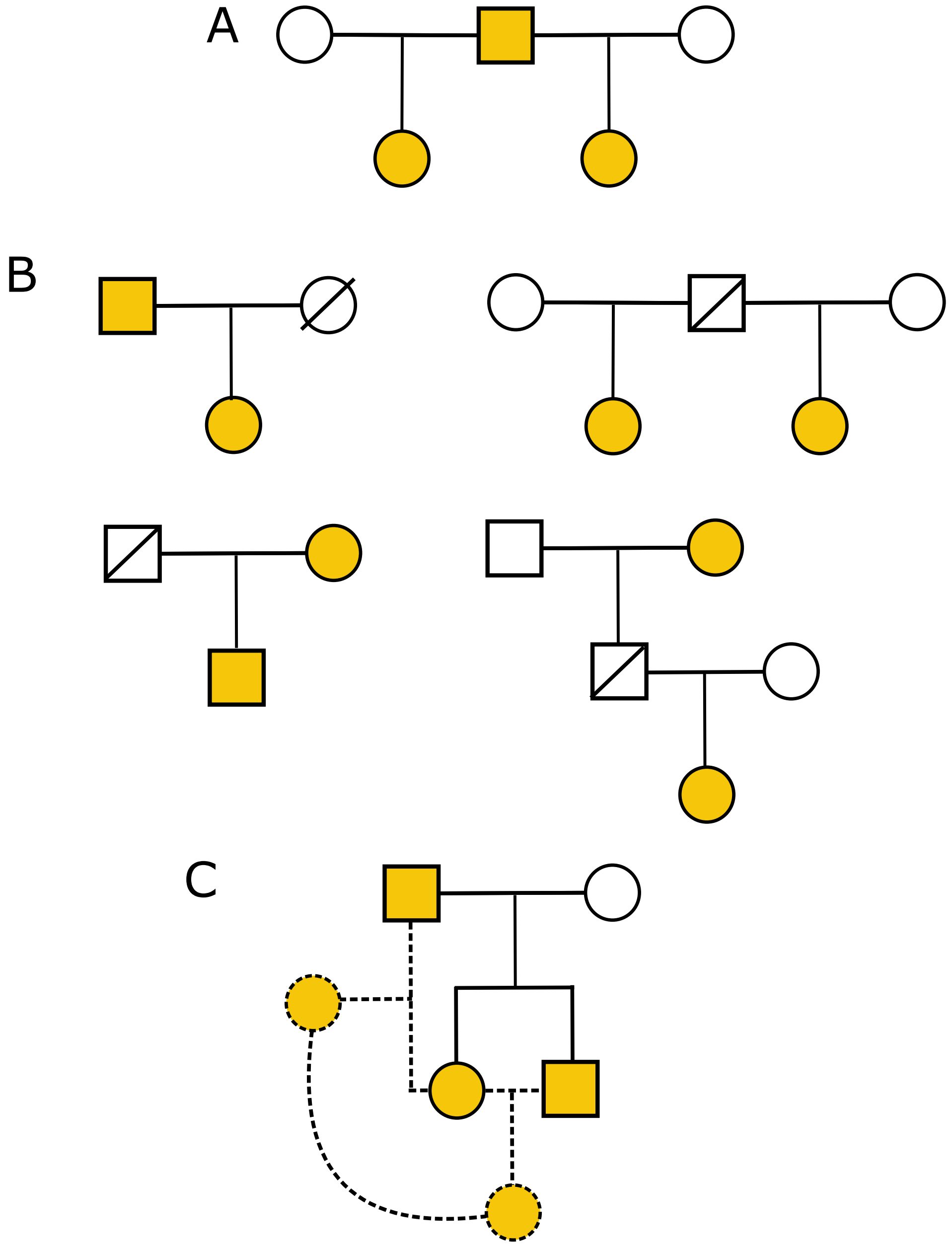

Fig. S1: Examples of kinship cases where X chromosomal markers may be informative: half-sisters (A), deficiency paternity test (B) and incest cases (C). The individuals whose genotype needs to be assessed to resolve the kinship case are in yellow.

### 2 Technical issues for X chromosome analyses

Some reasons for the general preference for autosomes in forensic identification and kinship analyses concern the fact that they may be analyzed regardless of the sex of the considered individuals. Moreover, autosomal multiplex assays are composed of about twenty markers, which are either located on different chromosomes or far from each other on the same chromosome, with their transmissions being considered independent events. In this way, the probabilities of the alternative hypotheses are built by multiplying together each genotype probability under the two scenarios. Conversely, when all markers are located on the same chromosome, such as in the case of X-STRs multiplexes, the informativeness given by considering just independent loci would not be sufficient to reach enough evidential weight. For this reason, recombination rates and haplotype frequencies should be considered in the biostatistical evaluation. Nevertheless, the paucity of dedicated software for the inference of recombination rates, as well as, the necessity to evaluate a large number of individuals for haplotype frequency estimation, introduce technical issues in X chromosome analyses.

#### 2.1 Linkage and Linkage Disequilibrium

Linkage and linkage disequilibrium come into play whenever physically close markers are considered, such as in X-STR multiplexes. Despite both being strongly related to chromosomal recombination, they indicate different events and require different biostatistical considerations (Tillmar *et al.,* 2017; Gomes *et al.,* 2020; Weir, 2019).

Linkage refers to the fact that physically close markers are more prone to be inherited together than physically separated or independent markers. On the other side, two markers are said to be unlinked if recombination is expected to occur in each meiosis, thus producing recombining gametic products half the time. In this case, the recombination rate would be 0.50 (Gomes *et al.,* 2020).

Linkage disequilibrium (LD) refers instead to alleles (and not loci): in particular, LD exists when alleles at different loci occur together, at a population-level, more or less often than what is expected by chance. The non-random association of alleles depends on population events, such as drift, selection, non-random mating or admixture. Moreover, the close physical location of the markers, as well as population stratification, increases LD. Being so dependent on population-specific characteristics which may vary across time, LD needs to be recomputed population by population and, even in the same human group, it is not stable, since recombination continuously breaks it (Gomes *et al.,* 2020).

However, both linkage and LD are unavoidable issues when performing X chromosomal analyses. Indeed, as inferrable from the length of the X chromosome, a maximum of four unlinked X-STRs may be analysed simultaneously (see the next section for more details). Concerning LD, X chromosomal markers exhibit higher LD values than autosomal ones, because X chromosomal recombinations occur just in female meiosis, whose mutational rate is also smaller (Schaffner, 2004). Moreover, haplotype frequencies cannot be inferred by simply multiplying together genotypic frequencies, because, due to LD, the condition of independence fails. Consequently, they should be computed by directly counting each haplotype across a population, meaning that larger databases are required.

##### 2.1.1 Linkage

Recombination rate studies, performed by evaluating either multi-generation families or large population databases (see below for more details), have generated maps of genetic distances, which are traditionally given in centiMorgans (cM).

Generally, it is assumed that no linkage between markers exists when the genetic distance is at least 50 cM. Several mapping functions can then be used to convert cM into recombination rates, such as Haldane and Kosambi (Haldane, 1919; Kosambi, 1943). For instance, as reported in Tillmar *et al.,* 2017, 50 cM corresponds to a recombination rate of 32% using Haldane's mapping function:

$$r = \frac{1 - e^{-\frac{2x}{100}}}{2} = \frac{1 - e^{-1}}{2} \approx 0.32$$

where $r$ is the recombination rate and $x$ is the genetic distance in cM. Interestingly, when using Haldane's or Kosambi's mapping functions, a recombination rate of 50% can be obtained just when markers are located at a distance of at least 200 cM. Thus, it is not advisable to assume an independent transmission for markers at less than 200 cM, unless otherwise inferred from informative families (Tillmar *et al.,* 2017).

Given that the X chromosome is roughly 155 Mb long, a maximum of 3-4 X-STRs separated by more than 50 cM will segregate independently. However, in order to increase the evidential weight, additional markers organized into "linkage groups" or "clusters" may be considered (Tillmar *et al.,* 2017).

The combination of groups of tightly linked X-STRs (linkage groups or clusters) into haplotypes, while increasing the evidential weight, also requires proper considerations in the biostatistical evaluation of kinship (Kling *et al.,* 2015b; Kling *et al.,* 2015a). The latest recommendations of the International Society for Forensic Genetics (ISFG) about the use of X-STRs in kinship analyses reported in Tillmar *et al.,* 2017 clearly indicate the precise knowledge of recombination rates between markers included in in-house and commercial X-chromosomal multiplex PCR assays as a prerequisite to unbiased estimates of kinship likelihood ratios (LRs):

**Recommendation 1:** *"Prior to using an X-chromosomal assay or commercial kit, markers should be evaluated to determine whether or not they are linked. Recombination rates should primarily be estimated from family studies or secondarily via mapping functions based on genetic distances. A recombination rate below 0.5 indicates linkage."*

**Recommendation 2:** *"Linkage should be accounted for when calculating LRs given that the X-chromosomal markers are linked and that linkage will have an impact on the final LR. This also includes accounting for recombination events within a cluster of X chromosomal markers, known as a linkage group."*

##### 2.2 Probability of mutation

The mutation rate of STRs ranges from $10^{-6}$ to $10^{-2}$ nucleotides per generation and is considerably higher than other kinds of polymorphisms, such as SNPs (Fan & Chu, 2007; Steely *et al.,* 2022). Mutations arise because the DNA replication process, despite being very accurate, can make mistakes inserting a wrong nucleotide or sometimes adding or eliminating others, at a rate of about 1 per every 100,000 nucleotides. Fortunately, most of these errors are fixed by DNA repair machines; however, in some cases, DNA mutations can escape those mechanisms, thus being passed down to the next generation. In particular, polymerase template slippage is the main molecular mechanism involved in STR mutations and mutations involving the loss or gain of just one repetition are predominant over those involving multiple repetitions (Gomes *et al.,* 2020).

In forensic practice, the estimation of mutation rates is crucial for the biostatistical evaluation of the evidential weight. Unfortunately, there is a general lack of research concerning the most commonly used STRs, mainly due to the fact that mutation rates can be assessed from allele-transmission in parent-child pedigrees, thus requiring a high number of specific family configurations (Steely *et al.,* 2022).

#### 3 The statistical framework of recombination rates inference

The physical location of X-STR along a single chromosome exhibiting peculiar inheritance mode and genetic recombination limited to females needs to be properly considered in the biostatistical evaluation of kinship, as also recommended by the International Society for Forensic Genetics (ISFG) (Tillmar *et al.,* 2017). However, despite being crucial for the unbiased formulation of the evidential weight, few biostatistical tools for the evaluation of recombination rates between adjacent forensic markers along the X chromosome are available today. Moreover, the main statistical approach based on pedigrees is unsuitable when more than 12-15 genetic markers are considered (Nothnagel *et al.,* 2012; Diegoli *et al.,* 2016). As a matter of fact, routine kinship analyses relying on X chromosomal markers are rarely performed with more than a handful of them, with the commercial kit Argus X-12 QS (consisting indeed of 12 STRs) being the most used in forensic practice. Nevertheless, the growing use of next-generation sequencing technologies in the forensic fields, with the possibility of combining thousands of markers together, requires the development of new biostatistical frameworks scalable to an arbitrary number of genetic markers.

The standard statistical approach for the estimation of recombination rates from pedigrees follows a comprehensive likelihood-based approach, while also allowing for meiotic mutations. This approach was developed using the programming language $R$ in 2012 by Nothnagel and colleagues (Nothnagel *et al.,* 2012) and represents, to the best of our knowledge the only available statistical approach to evaluate pedigree-based recombination rates. However, it computes the likelihood of kinship by taking into account all possible recombinations within the maternal haplotype, thus exhibiting an exponential time complexity. The time complexity of an algorithm is expressed using the $O$ notation and indicates how quickly its run time increases relative to the input $n$. In the work of Nothnagel and colleagues, the number of computations, which is a proxy of the overall necessary time to complete a task, doubles with each item added to the input ($n$) and can be written as $O(2^n)$. Exponential time complexity is common in those situations where every possible data combination (in this specific case, every possible recombination within the maternal haplotype) needs to be explored in order to get a clue about the best solutions. For this reason, the underlying algorithm complexity prevents the estimation of recombination rates for more than 12 STRs (Nothnagel *et al.,* 2012).

A computation update from the same research group came out in 2016 (Diegoli *et al.,* 2016). In this work, they modified the previous approach by writing the underlying code in C++ and introducing code parallelization in order to allow the computations to be performed on a high-performance computing cluster. Nevertheless, even this approach is reported to be unsuitable when panels of more than 15 X-STRs are considered (Diegoli *et al.,* 2016).

#### 3.1 Likelihood formulation

The statistical approach for the estimation of recombination rates carried on by Nothnagel and colleagues relied on comprehensive likelihood calculations, which also took into account the possibility of single-step mutations (see the Materials and Methods section in the main text). In particular, fractional mutations or those involving more than one repeat were considered too unlikely and the entire family was discarded from the analyses. We report below the thorough explanation of the statistical approach used in Nothnagel *et al.,* 2012 and Diegoli *et al.,* 2016, using the original notation (conversely, in the Materials and Methods section of the main text, we use a slightly modified but more straightforward notation—in our opinion).

For both type I and type II families, the likelihood of a family genotype is formulated as follows: given $n$ X-STRs for which the genomic position and so the physical order is known, let $(\theta_1, ..., \theta_{n-1})$ be the recombination rates between adjacent markers. $\mu$ indicates instead the one-step mutation rate which is uniform and symmetric, meaning that a single mutation rate which applies to all X-STRs is estimated.

An "inheritance vector" can be built for each mother-son pair: $V \in \{1, 2\}^n$ is the inheritance vector, where each element $V(i)$ can assume values 1 or 2 depending on the grandparental origin of the allele at the $i$-th STR (for instance, $V(i)$ = 1 for grandpaternal, $V(i)$ = 2 for grandmaternal; Figure S2).

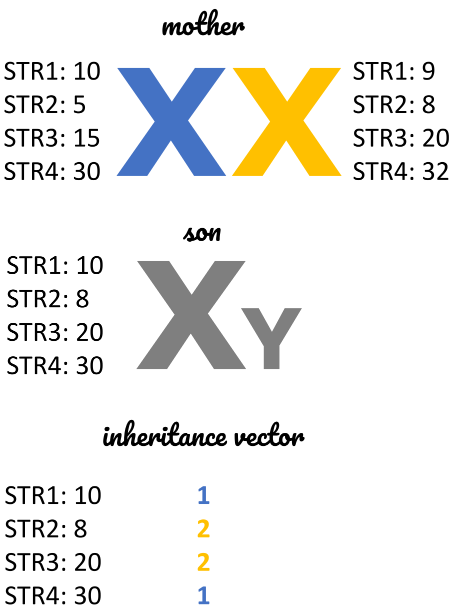

Fig. S2: One of the possible inheritance vectors for a given mother-son pair.

The conditional likelihood of the genotype $g_s(i)_{i=1..n}$ of the son, given the phased maternal genotype $g_m(i, j)_{i=1..n, j=1..2}$ and an inheritance vector $V$, is:

$$L(g_s \mid g_m, \theta_1, ..., \theta_{n-1}, \mu, V) = \prod_{i=1}^{n-1} [\theta_i + (1 - 2\Theta_i) \cdot 1_{V(i)=V(i+1)}] \times \prod_{i=1}^{n} [\mu + (1 - 2\mu) \cdot 1_{g_m(i, V(i))=g_s(i)}] \tag{1}$$

The first part of the formula refers to the recombination, while the second indicates the mutation events. Starting from the end of the recombination part, there is the *indicator function* $1_{V(i)=V(i+1)}$, which takes value 1 when the condition is satisfied or 0 when it is not. This function checks whether, within the inheritance vector $V$ the $i^{th}$ position is equal to the $i^{th}+1$, thus indicating that no recombination has happened. In this case, by solving the recombination part, we obtain $1-\theta$: if $\theta$ is the probability of recombination, $1-\Theta$ is simply the probability that recombination did not happen. On the contrary, if the maternal haplotype changes between marker $i$ and $i+1$, this means that recombination did happen: in this case, the indicator function takes the 0 value and the only term left is $\Theta$, which indeed is the probability of recombination. In the second part, the indicator function $1_{g_m(i,V(i))=g_s(i)}$ is 1 if the mother genotype on the haplotype inherited by her son (as indicated by the inheritance vector $V(i)$) is identical to the son genotype, meaning that no mutation occurred. Thus, we obtain $1-2\mu$ with no mutations and $\mu$ when a mutation occurred. Notably, we introduced a slight modification of the original formula (1), as in Nothnagel *et al.,* 2012: if there is a fractionary mutation or one involving more than 1 step, the likelihood for that particular inheritance vector is 0.

While the first part iterates from 1 to $n-1$, the second goes from 1 to $n$ because recombination happens *between* markers, while mutation may occur at *each* marker. Finally, the two parts are multiplied together to obtain the likelihood of a son given a particular $V$.

Then, summing the term in 1 over all $2^n$ possible inheritance vectors, given the phased maternal genotype alone, gives the conditional likelihood of the genotype of the son:

$$L(g_s \mid g_m, \Theta_1, ..., \Theta_{n-1}, \mu) = \sum_{j=1}^{2^n} L(g_s \mid g_m, \Theta_1, ..., \Theta_n, \mu, V_j) \tag{2}$$

As already mentioned, having available the grandpaternal genotypes in type I families, allows the phasing of the maternal genotypes. In this way, the likelihood of the whole family is simply the product of the son-specific likelihoods as given in (2) (this happens because mother's recombinations are independent events, i.e., mother's X chromosome recombined independently for each son). Indeed, in type I families, one child is enough to provide information on recombination, while, in type II families, at least two children are needed. However, in type II families, the maternal phase is usually unknown, so a certain degree of uncertainty needs to be taken into account for likelihood calculation by summing the aforementioned likelihood products over all $2^n$ possible maternal phases.

In the end, the likelihood of the total data is obtained by multiplying all family-specific likelihoods (Nothnagel *et al.,* 2012) (again, for the independence among families).

#### 3.2 Likelihood maximization

Since likelihood functions can not be directly used to compute the recombination rates, Nothnagel and colleagues used the expectation-maximization method (EM), which allows to "reverse" the likelihood functions. The basic idea is to find the recombination rates that best fit the observed dataset, i.e., that maximize its likelihood.

The procedure starts from an initial approximation of the recombination rates that are used to compute the initial global likelihood. Then, the recombination rates are modified slightly and the likelihood is recomputed. If it increases, then the modified rates that better fit the observation are retained, otherwise, they are discarded and a new iteration with other modified rates carries on. This process is repeated again, producing a series of improving estimates of the recombination rates until the likelihood stops increasing. That is the final estimate. This process is generally not perfect: the recombination rates it produces may not be the absolute best and this depends a lot on the initial approximation of the recombination rates. In order to control for this, the EM is run multiple times with different initial approximations.

This procedure was implemented by Nothnagel and colleagues in R v 2.13.0, using in-house scripts, which are available from the authors upon request. In detail, likelihood maximisation was performed using the *optim* function and the methods "L-BFGS-B" which allows box constraints, that is each variable is given a lower and upper bound, with the initially provided value satisfying the constraints (Byrd *et al.,* 1995). In this case, recombination rates were constrained to be among $10^{-8}$ and 0.5, while a mutation rate $\mu$ of 0.001 was employed for all markers. This number came from estimates for X-STRs (Liu *et al.,* 2011; Castañeda *et al.,* 2012) and was confirmed by the very same findings of Nothnagel *et al.,* 2012: indeed, they found 8 mutations in 10,290 informative meioses, which corresponds to $7.8 \times 10^{-4}$.

#### 3.3 The dynamic programming implementations

Our fastest implementations are based on "Dynamic Programming", a computer programming technique useful when dealing with problems that have overlapping sub-problems. Basically, the main concept behind dynamic programming is to divide the problem into smaller sub-problems: in case of overlaps among sub-problems, their solutions can be stored, in order to avoid repeating the same computations. A classical example of dynamic programming usefulness is the Fibonacci sequence, where each number is the sum of the previous two. The Fibonacci sequence is:

$0, 1, 1, 2, 3, 5, 8, 13, 21, 34, 55, ...$

Finding a number of the Fibonacci sequence starting from its $i^{th}$ position within the sequence can be solved using a recursive algorithm as follows:

```
def fib_val(n):
    if n == 0:
```

```
        return 0
    elif n == 1:
        return 1
    else:
        return fib_val(n-1) + fib_val(n-2)
```

In this way, fib_val(5) = 5. However, in order to compute the f(5) (Fibonacci value for the $5^{th}$ position), we have to compute, for instance, f(2) 3 times.

$$\begin{aligned} f(5) &= f(4) + f(3) \\ &= f(3) + f(2) + f(2) + f(1) \\ &= f(2) + f(1) + f(2) + f(2) + f(1) \end{aligned} \tag{3}$$

The dynamic programming solution for this problem consists in computing and memorizing this value just one time.

We report below the dynamic programming-based Python function for the inference of type I family likelihood.

```
def compute_phased_family_likelihood_dyn_loop(mother,
   maternal_haplotypes, recombination_rates, mutation_rates):
    """Simple dynamic programming implementation of phased (type I)
   family likelihood.

    The dynamic programming algorithm is implemented explicitly with
   nested loops.
    This function can be compiled with numba if available for a big
   speedup.
    """

    mode = 'SNP' if mother.dtype == '<U1' else 'STR'
    lh = 1.0
    m = numpy.zeros(shape=mother.shape)
    for hap in maternal_haplotypes:
        for pos in range(mother.shape[0]):
            for r in range(2):
                # compute mutation probability
                if mode == 'SNP':
                    mut_step = mother[pos, r] != hap[pos]
                elif mode == 'STR':
                    mut_step = abs(mother[pos, r] - hap[pos])
                if mut_step == 0:
                    mutp = 1 - mutation_rates[pos]
                elif mut_step == 1:
                    mutp = mutation_rates[pos]
                else:
                    mutp = 0
                # compute recombination probability
                m[pos, r] = 0.5*mutp if pos == 0 else mutp*(
```

```
                    m[pos - 1, r]*(1 - recombination_rates[pos - 1])
 +
                    m[pos - 1, 1 - r]*recombination_rates[pos - 1]
            )
        lh *= numpy.sum(m[-1])
    return lh
```
Listing 1: The *Dynamic* implementation.

If available, this implementation may be combined with Numba, a "compiler" for Python which works by translating Python functions to optimized machine code, thus approaching the speeds of C or FORTRAN (Lam *et al.,* 2015). Python is an "interpreted" language, meaning that the code is not directly executed, but interpreted by another program. Compiled languages are instead compiled (and not interpreted): when the code is compiled, it is expressed into machine instructions, which are undecipherable by humans. Examples of interpreted languages are Python, R, Perl and so on, while C++ is a compiled language. Given that interpreted languages need another program to interpret the code, they tend to be slower and less efficient. Hence, Numba can be used to speed up the computation, by simply wrapping the chosen function into *numba.njit*.

### 4 Recombulator-X

Recombulator-X is a Python module and a command line tool for computing the recombination and mutation rates between X-STRs starting from pedigree data.

The code of the tool is freely available on GitHub. Full documentation can be found on the dedicated website:

**GitHub**: https://github.com/serena-aneli/recombulator-x
**Website**: https://serena-aneli.github.io/recombulator-x/

#### 4.1 Installation

The tool can be easily installed via pip, the package installer for Python, as follows:

```
pip install recombulator-x
```

In order to properly work, Recombulator-X needs the following Python modules and versions: numpy>=1.14, pandas>=0.23, scipy>=1.0, networkx>=2.0. Notably, it is not mandatory to install Numba: if Numba is not installed, just the implementations which do not use Numba will be available.

#### 4.2 Usage

##### 4.2.1 Input files

Recombulator-X uses the PED format based on PLINK pedigree files as input (Chang *et al.,* 2015). The PED file format stores sample pedigree information (i.e., the familial relationships between samples) and the genotypes (Table S1). In particular, the first 6 mandatory columns contain:

- Family ID
- Individual ID
- Paternal ID
- Maternal ID
- Sex
- Phenotype

The "Sex" field may be coded as: 1=male/ 2=female; XY=male/ XX=female; M=male/ F=female; MALE=male/ FEMALE=female. In the case of STR markers, the Amelogenin marker (which indicates the sex) can be stored within this column. Other kinds of polymorphisms may be coded as single-letters (SNPs), but generic strings are accepted in the case of more complex polymorphisms like INDELs.

The "Phenotype" field comes from medical research tradition. In non-medical applications, it may be -9 which means "unknown".

From the 7th column on, there are the markers' genotypes (two columns for a genetic marker, each of the two storing an allele). In the case of STRs, the columns contain numbers, which correspond to the STR repeats or "0" when missing.

**Recommendation** : Genetic markers (from the 7th column on) must be provided according to their physical genomic position. Indeed, the algorithm will infer the recombination rate between A1 and A2, A2 and A3 and so on.

Here is a family (each row is an individual):

| FID | IID | PAT | MAT | SEX | PHENO | STR-A1 | STR-A2 | STR-A1 | STR-A2 | STR-A1 | STR-A2 |
|---|---|---|---|---|---|---|---|---|---|---|---|
| FAM_I | GRANDFATHER | 0 | 0 | 1 | -1 | 12 | 0 | 29 | 0 | 39 | 0 |
| FAM_I | MOTHER | GRANDFATHER | 0 | 2 | -1 | 12 | 16 | 27 | 29 | 34 | 39 |
| FAM_I | SON_1 | 0 | MOTHER | 1 | -1 | 12 | 0 | 29 | 0 | 34 | 0 |
| FAM_I | FATHER_1 | 0 | 0 | 1 | -1 | 14 | 0 | 21 | 0 | 37 | 0 |
| FAM_I | DAUGHTER_1 | FATHER_1 | MOTHER | 2 | -1 | 14 | 16 | 21 | 27 | 34 | 37 |
| FAM_I | FATHER_2 | 0 | 0 | 1 | -1 | 18 | 0 | 25 | 0 | 36 | 0 |
| FAM_I | DAUGHTER_2 | FATHER_2 | MOTHER | 2 | -1 | 12 | 18 | 25 | 29 | 36 | 39 |

Table S1: Some lines from a simulated PED file (each row is an individual).

#### 4.3 Assumptions and workflow

Before delving into the features of the tool, it may be worth mentioning its main assumptions:

- Males must be haploid for all the markers: given that our tool is designed for X-chromosomal markers, males have just one copy of the X-chromosome.
- For the current version of Recombulator-X, markers must be short tandem repeats (STR).
- Unit mutations: STR fractionary mutations are not allowed and mutation of more than one repeat are assumed to have zero probability
- Genetic markers on the PED files must be provided according to their physical genomic position.

The program can be used both as a Python module and a command-line tool. We will report below the workflow of Recombulator-X as well as some indications about its usage.

###### 4.3.1 Pedigree preprocessing

The initial steps of Recombulator-X consist in reading the PED file and identifying the informative families for the estimation of recombination rates (function *ped2graph*). Moreover, using the Python package *networkx*, we also added an interesting feature: indeed, starting from the relatedness reported within the columns “PAT” and “MAT” of the PED file, the tool can reconstruct the family graph.

After the input file reading, the function *ped2graph* performs also some checks on the families: for instance, the columns need to be even (6 initial columns, plus two columns per marker). Then, each individual is added as a node to an empty graph, while the edges connect the related individuals (Figure S3).

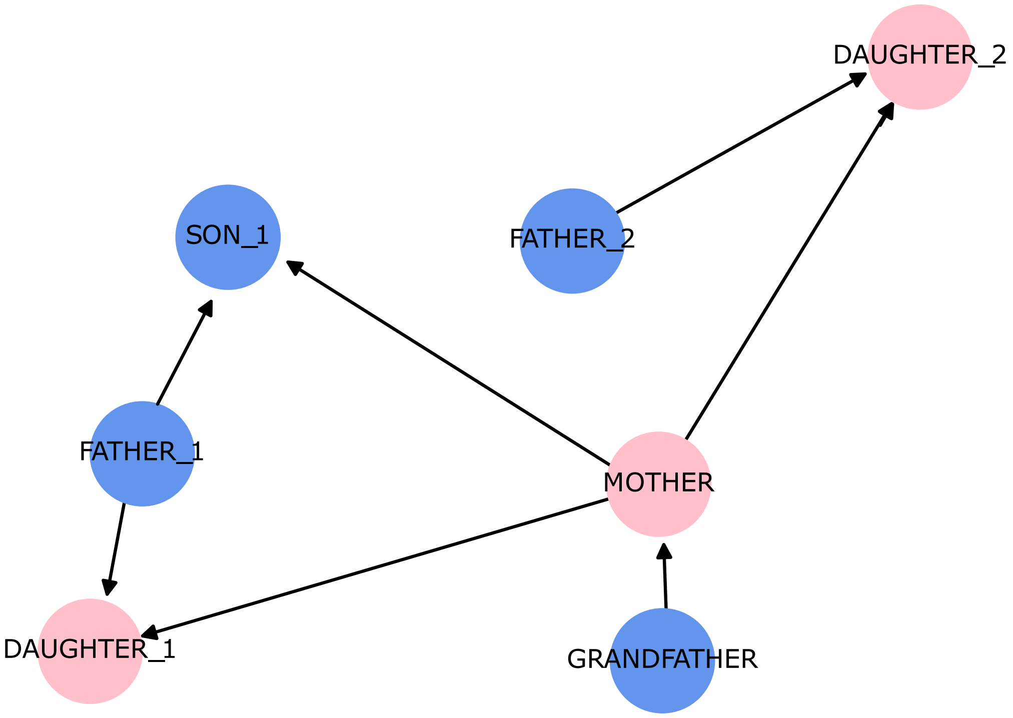

Fig. S3: Graph of a simulated family.

As a second phase, we also added a dedicated function to check the consistency of each family graph and raise errors whenever necessary (function *check˙family˙graph*). For instance, an error is raised when more than two parents or same-sex parents are present in the same family. Unconnected individuals are also flagged.

At this point, the tool will count how many informative type I and type II families (function *extract˙informative˙subfamilies*). For recombination, informative subfamilies are either those with:

- a phased mother and at least one son or phased daughter called type I families

- an unphased mother and at least two between sons and phased daughters, called type II families

Note that these families are informative also for mutation. The following lines perform specifically this task, thus representing one the most important parts of the tool.

Notably, females can be phased when their father is available: in this way, they will be virtually transformed into males, thus being allowed to take part in informative families. The function *phase˙daughter* following function performs the phasing of females, whenever their father is present.

###### 4.3.2 Likelihood estimation

After the family preprocessing, Recombulator-X will perform the necessary steps to infer recombination and mutation rates:

1. computation of the likelihood of a son given his mother's genotypes, inheritance vector, recombination and mutation rate;
2. we sum the likelihoods computed over all possible inheritance vectors (these are disjoint events since just one inheritance vector will be true);
3. we multiply the likelihoods of all sons within a family in order to obtain the likelihood of the family;
4. for type II families, we do not know the mother phasing, thus, we repeat the previous steps computing the family likelihood for all possible mother phasings and sum them (as disjoint events);
5. we multiply the likelihoods of all families in order to obtain the likelihood of the entire dataset;
6. actually, we are not so interested in the likelihoods themselves, but in their parameters, which are the recombination and mutation rates. Hence, using EM, we find the parameters maximizing the overall likelihood.

The user can decide which of the likelihood implementations developed in Recombulator-X will be used to perform the first three steps:

- For loop implementation of the *direct* algorithm (direct-loop);
- NumPy implementation of the *direct* algorithm (direct-numpy);
- *Dynamic programming* algorithm (dynamic );
- A compiled version of the for loop implementation of the *dynamic programming* algorithm (dynamic-numba).

##### 4.3.3 Practical usage

Recombulator-X can be used both as a Python module and a command-line tool. File *Estimation Example.ipynb* within the GitHub repository reports a detailed notebook for the Python module. Additional information is also present in the online documentation.

The estimation of recombination and mutation rates can be launched with the following line:

```
# estimate_rates(families, starting_recombination_rates=0.05,
    starting_mutation_rates=0.001, estimate_mutation_rates='no',
    implementation='auto')
est_recomb_rates, est_mut_rates = recombulatorx.estimate_rates(
    processed_families, 0.1, 0.1, estimate_mutation_rates='all')
```

The function *estimate˙rates* estimates recombination and mutation rates from a set of families and takes the following parameters: the families, the initial recombination rate, the initial mutation rate, which mutation rate needs to be estimated (*no*: no mutation rate estimation, *one*: just one mutation rate for all markers, *all*: a mutation rate for each marker), the type of implementation (the default implementation is the one using dynamic programming).

An example of output generated by function *estimate˙rates* in Python is:

```
(array([0.03874299, 0.32869992, 0.01459788, 0.19265765, 0.01016452]),
 array([1.00000000e-08, 1.00000000e-08, 1.23511804e-01, 2.10614659e
    -02, 2.24679981e-03, 1.00000000e-08]))
```

where, the first array ($n-1$ long) stores the recombination rates, while the second ($n$ long) contains the mutation rates estimated for simulated families and six X-STRs.

The command line interface for Recombulator-X was created using the Python library *argparse*. The tool has the following parameters:

```
usage: recombulator-x [-h] [--mutation-rates MUT-RATE [MUT-RATE ...]]
                      [--estimate-mutation-rates {no,one,all}]
                      PED

Estimate recombination and mutation rates.

positional arguments:
  PED                   path to ped file

optional arguments:
  -h, --help            show this help message and exit
  --mutation-rates MUT-RATE [MUT-RATE ...]
                        mutation rates used in the estimation, either
                        as fixed or as starting point in the
                        optimization depending on the value of the
                        --estimate-mutation-rates option. If not
                        given the rates are set to 0.001 for all
                        markers
```

```
 --estimate-mutation-rates {no,one,all}
                        controls the estimation of the mutation
                        rates. With "no" the mutation rates are not
                        estimated, with "one" the same rate is
                        estimated for all markers, with "all" a
                        separate estimation rate is estimated for
                        each marker. Defaults to "no"
```

Its base usage consists in estimating just recombination rate and using the default single value for mutation rate (0.001). This task can be performed as follows:

```
recombulator-x ped_path
```

However, we may also decide to estimate also mutation rates. In particular, adding *–estimate-mutation-rates all*, the tool will compute a mutation value for each marker.

```
recombulator-x ped_path --estimate-mutation-rates all
```

As an additional feature, Recombulator-X can also simulate pedigrees and their genotypes, a useful feature when testing new approaches (function *generate˙complex˙families*).
